# MyVivarium: A cloud-based lab animal colony management application with realtime ambient sensing

**DOI:** 10.1101/2024.08.10.607395

**Authors:** Robinson Vidva, Mir Abbas Raza, Jaswant Prabhakaran, Ayesha Sheikh, Alaina Sharp, Hayden Ott, Amelia Moore, Christopher Fleisher, Hailey Netherton, Evan Goldstein, Pothitos M. Pitychoutis, Tam V. Nguyen, Aaron Sathyanesan

## Abstract

Management of research-animal colonies is vital for lab productivity in preclinical research. Labs often rely on inefficient paper-based or spreadsheet-based methods to manage animal colonies. Dedicated software-based solutions are generally expensive and many lack remote access. Currently available open-source alternatives are difficult to implement and deploy. These solutions also do not have the capability to track ambient variables that affect colony well-being. We built *MyVivarium*, an open-source database management web application to address these gaps. MyVivarium can be easily deployed to the cloud and sustained for a cost comparable to starting and maintaining a lab website. Using MyVivarium, lab members can collaboratively track individual animals within a database. Physical identities of cages map onto the database using QR codes, enabling quick and easy record-keeping on mobile devices. Lab administrators can assign tasks to users with reminders for experiments or cage maintenance. Finally, we designed a low-cost system to sense ambient humidity, temperature, vivarium worker activity, and room illuminance. These data are then sent to MyVivarium in realtime providing information relevant to colony well-being. Taken together, MyVivarium is a novel, open-source, cloud-based application template that provides a low-cost, simple, and efficient way to digitally manage research-animal colonies.

## INTRODUCTION

Animal colony management is a routine task performed by many laboratories that use model systems to answer biological questions. Labs that use rodent models often are responsible for husbandry and the well-being of animal subjects under their purview. This can result in the day-to-day management of hundreds of animal cages per lab, making cage information management a key problem. Large mouse repositories use dedicated software to increase efficiency in mouse colony management. Yet, barriers remain in the average academic lab choosing a dedicated solution [1].

Most makeshift solutions for mouse colony management are *Google Sheets* or Microsoft Excel spreadsheets. Lab members log colony updates to these spreadsheets in a semi-collaborative manner [2]. During a typical colony inspection, a researcher would visit the vivarium, carrying along a paper copy of the colony cage list. Following routine updates to the paper list, this information is then usually updated in the spreadsheet database at a later time. Information directly related to a particular strain, such as genotype visualized on a gel image, is usually maintained separately in a lab notebook. Many labs have adopted this method of managing mouse colony management, however, it is cumbersome and has multiple points of failure due to its reliance on disparate tools and the lack of an integrated framework.

Rodent strains used as disease models in preclinical research often have genetic modifications that make them more sensitive to changes in ambient temperature, relative humidity, and light [3–6]. In addition, human presence may also negatively affect genetically modified mouse strains. Changes in vivarium ambient factors can thus negatively impact colony breeding efficiency and wellbeing. Commercial solutions to monitor ambient factors in rodent colonies do exist, however, these are significantly cost-prohibitive [7]. A low-cost colony management tool that can also track alterations in ambient environmental factors is an immediate need for the research community.

The advent of the modern web has brought with it ever-improving tools for rapid and remote management of scientific data [8,9]. Early career researchers are increasingly conversant with designing websites for sharing their research programs [10]. We reasoned that if the setup and use of a rodent colony management system were as easy as launching and maintaining a lab website, more labs would adopt such a system.

Recently, some open-source database applications for rodent colony management solutions have been introduced [7,11]. However, these systems were primarily implemented to run on local servers and networks, limiting their access, especially in situations of contingency [12,13]. While these solutions can be moved to the internet, it is not trivial to do this securely and seamlessly. For many biologists with limited knowledge of web development, this is especially difficult to carry out. Furthermore, no open-source lab animal colony management tool has integrated ambient sensing. In this manuscript, we describe the design of *MyVivarium* – an open-source application framework for lab animal colony management with cloud-based access as the primary starting point. We built MyVivarium using a PHP-based technology stack running on a managed cloud-hosted virtual private server. Importantly, we prioritized cost by using an inexpensive server option (∼$6 USD per month per lab for 1 GB RAM, 25 GB SSD, 1 TB Transfer, 1 Core Processor). MyVivarium enables all the necessary standard operating procedures for vivarium maintenance, including cage and mouse record maintenance, with the ability to generate cage cards with unique QR codes for easy mobile access and tracking. In order to efficiently track experiments linked to mice in specific cages, tasks and reminders can be set up with designated researchers assigned to these tasks. Finally, we describe the assembly and integration of automated low-cost Raspberry Pi IoT (RPi-IoT) ambient sensors that continuously stream temperature, humidity, illuminance, and vivarium room activity data to MyVivarium in realtime. Fluctuations in ambient environmental factors based on user-defined criteria initiate automated alerts to designated personnel. MyVivarium is thus a complete open-source solution for low-cost, cloud-based lab animal colony management suitable for academic research labs as well as vivarium management staff.

## RESULTS

We aimed to create a simple and efficient interface for cloud-based cross-device-compatible mouse colony management. To this end, we built MyVivarium using modern web technologies for both client-side and server-side^1^ operations (**Figure 1**). We used the scripting language *PHP* for server-side scripting and *MySQL* for database management. On the client-side, we utilized *HTML*, *CSS*, and *JavaScript*, including the *Bootstrap* framework for responsive design and user interface components. We used the *Composer* tool for dependency management, incorporating libraries such as *PHPMailer* for email functionality and *Dotenv* for environment variable management.

**Figure 1.**
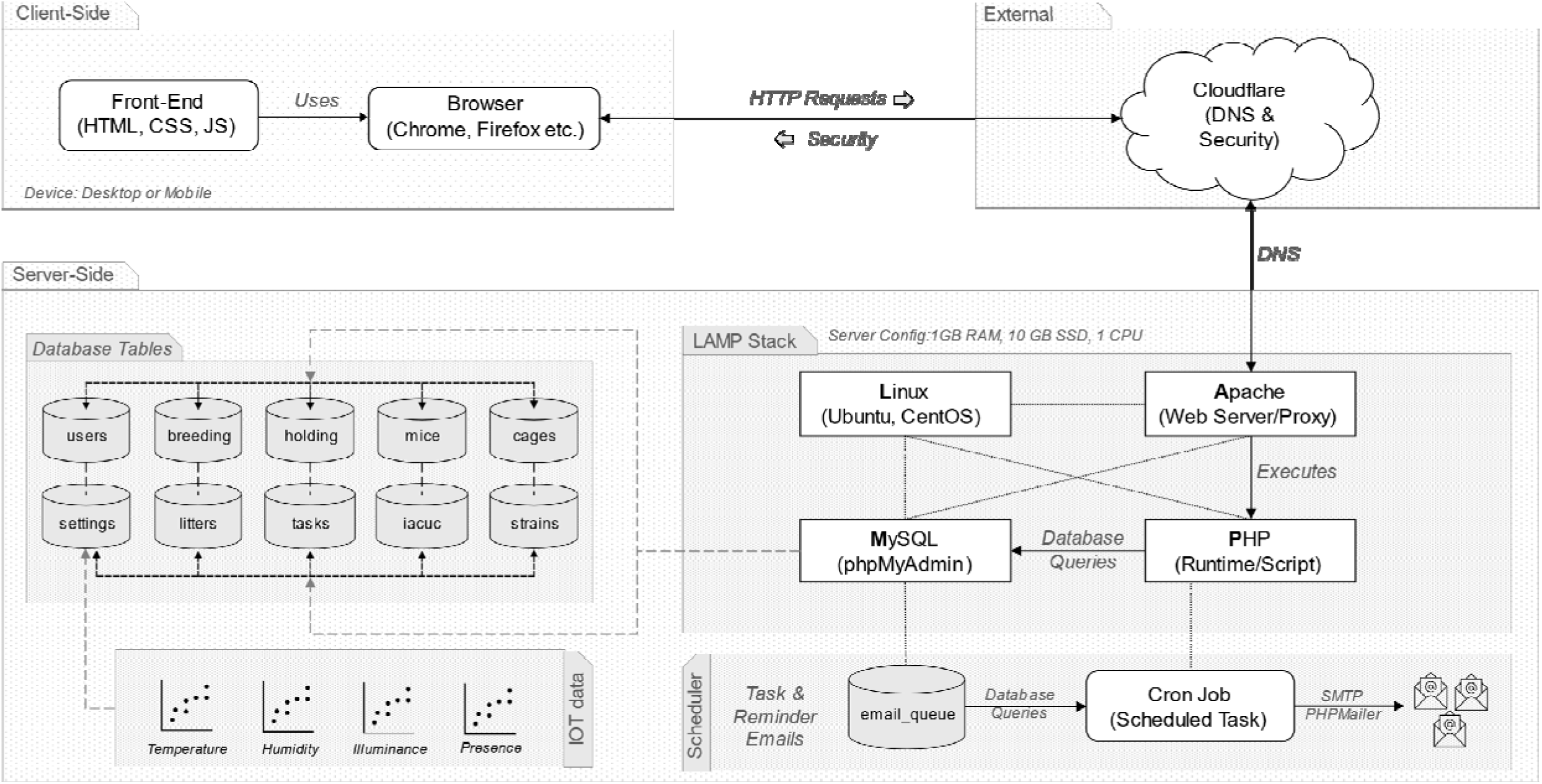
MyVivarium System Architecture Overview. MyVivarium system architecture consists of three main components: the client side, the server side, and external services. Each component interacts with the others to provide a robust and secure web application for managing vivarium operations. *Client-side:* MyVivarium’s front-end is built using standard web technologies, including HTML, CSS, and JavaScript, ensuring that the user interface is interactive and responsive. Users can interact with the application through web browsers such as Chrome and Firefox with cross-device compatibility. *External services:* Cloudflare provides DNS services and security features, including DDoS protection, to enhance the reliability and security of the MyVivarium application. *Server-Side*: The backend is built on a LAMP stack, consisting of Linux (operating system), Apache (webserver), MySQL (database), and PHP (scripting language). The database includes tables such as `users`, `breeding`, `holding`, `mouse`, `files`, `settings`, `litter`, `tasks`, `notes`, and `strains`, which store various types of data required for the application’s functionality. *IoT Data*: The system integrates IoT sensor data to monitor ambient parameters such as temperature, humidity, illuminance, and presence. *Scheduler:* A scheduler (cron job) handles tasks such as sending reminder emails utilizing the `email_queue` for processing. Cron jobs automate database queries and sending emails through SMTP using PHPMailer.

Key features of MyVivarium include user registration and profile management, lab and cage management, as well as real-time ambient sensing with internet-of-things (IoT) sensors (**Figure 2**). The application supports secure session handling, cross-site request forgery (CSRF) protection, and role-based access controls. The user management functionality includes features for registration, login, profile management, email verification, and password reset via email. Lab and cage management modules enable users to add, view, edit, and delete breeding and holding cages. Each module provides detailed information about the cages, including associated litter and mouse data. Additionally, users can manage strains by adding, editing, and deleting strain records. We have shared all necessary code and a how-to guide in a public repository (https://github.com/myvivarium/MyVivarium) for users to get their specific MyVivarium web applications set up according to their particular needs and requirements. We have set up a website where potential users can demo the different functionalities within MyVivarium (https://demo.myvivarium.online/). We have also launched a separate website to introduce potential users to MyVivarium: https://myvivarium.github.io/.

**Figure 2.**
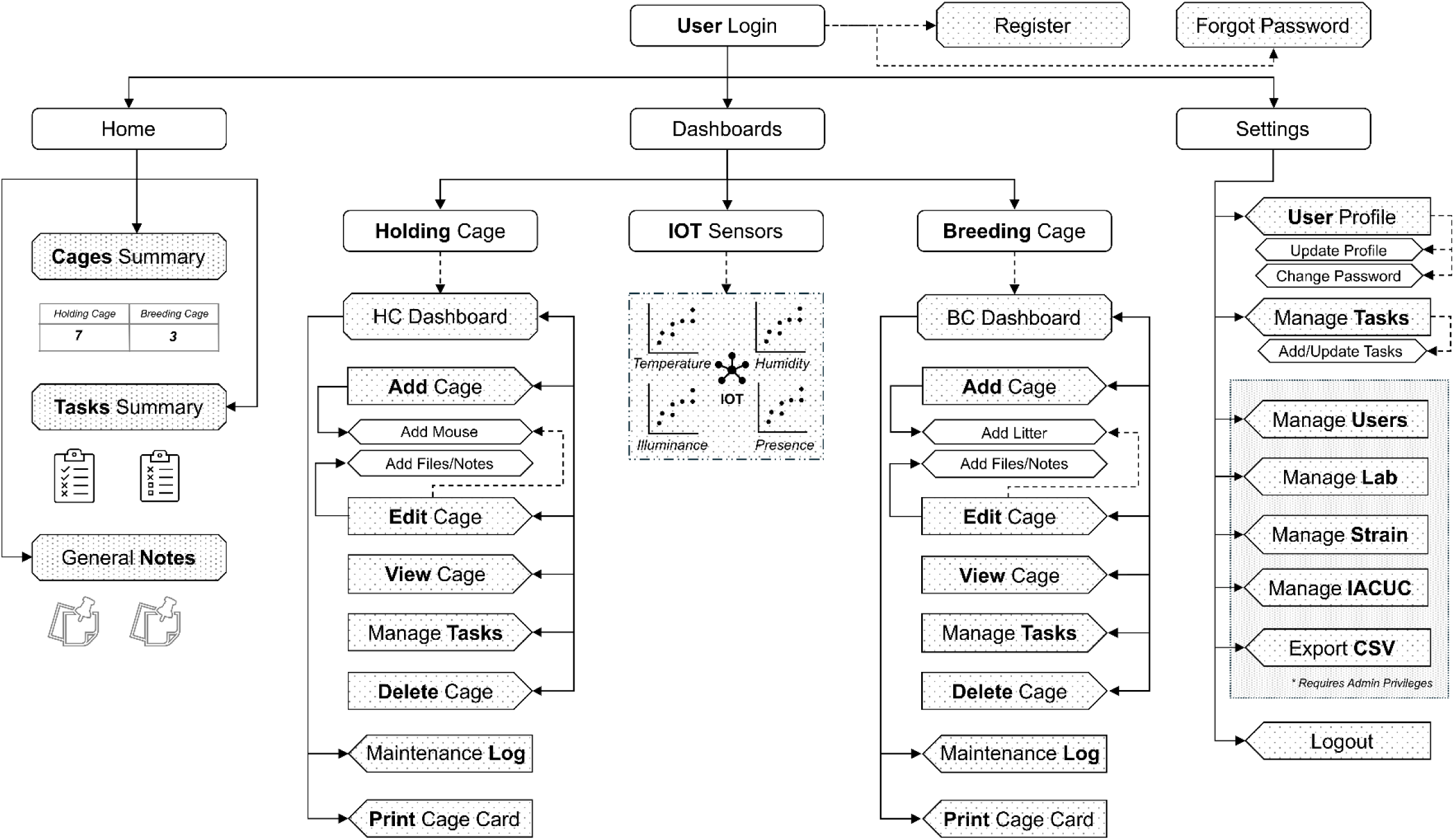
MyVivarium web application navigation overview. The MyVivarium web application navigation is intuitive, structured into “Home”, “Dashboards”, and “Settings”. The “Home” section includes “Cages Summary”, “Tasks Summary”, and “General Notes”. The “User Login” provides links for registration and password recovery. “Dashboards” gives access to holding cage and breeding cage management, including adding, editing, viewing, and deleting cages, printing cage cards, maintenance logs, and realtime ambient sensing via the IoT sensor datastream. “Settings” includes user profile management, task management, and admin options such as managing users, lab details, mouse strains, IACUC protocols, and data export in CSV format. The Logout option allows users to log out securely. This structure ensures easy access to all functionalities, with specific options available based on user roles and permissions.

Cage, mouse, and litter management forms the core of MyVivarium (**Supplemental Figure 1**). We designed the database architecture using active feedback from vivarium workers over the course of the development of MyVivarium. The MyVivarium database includes the `cages` table, which forms a core part of the architecture and contains essential information about each cage, including the cage ID, the name of the principal investigator, the number of mice, and any specific remarks linked to a cage (**Supplemental Figure 1**). Linked to this core table is the `users` table, which holds necessary information about users, including their role within the lab, position or designation, and MyVivarium access approval status. This ensures that each cage can be associated with specific users, facilitating proper access and management of responsibilities. The `holding` table stores essential information about individual mice in holding cages, including their strain, date of birth, sex, and parent cage. This table ensures that all necessary details about each holding cage are recorded systematically. The `mouse` table stores detailed information about individual mice within the holding cages. The database architecture for breeding cages in MyVivarium includes the `breeding` table, which contains information about each breeding pair, including their source cage IDs, type of cross, male and female IDs, and their dates of birth. Linked to this table is the `litters` table, which records details about offspring, such as the number of pups per litter, the number of surviving pups, sex, and any other relevant remarks. This setup enables monitoring and analysis of breeding outcomes per cage. The `strains` table contains information about various strains of mice, including strain IDs, names, and other identifiers, enabling tracking of different genetic lines. In order to link documentation related to the institutional animal care and use committee (IACUC) to specific strains, we also included an `iacuc` table. In comparison to previous colony management tools, our IACUC functionality enables better integration of IACUC-related tasks to help with the preparation of compliance and animal usage reports. Additional tables, including `files` and `notes`, support the documentation and communication needs within MyVivarium. The `files` table stores metadata about uploaded documents, while the `notes` table enables users to record observations and updates related to specific cages or mice. The `tasks` table manages tasks and assignments, tracks their status, and enables increased efficiency for time-sensitive vivarium tasks in the range of days-to-months. For more day-to-day tasks such as cage-changes and regular maintenance, we included a `maintenance` table to enable quick recording of maintenance logs. Finally, the schema also includes the `cage_iacuc` and `cage_users` tables to establish relationships between cages, IACUC protocols, and users. Together, these tables create a robust framework for vivarium workers to manage and track details of holding cages and the individual mice within them, as well as breeding cages and the breeders and litters within them.

From a user-experience perspective, once approved users are logged in to their lab’s MyVivarium, holding and breeding cages are accessible in the web application via the “Dashboards” menu (**Supplemental videos 1-4** [desktop]**, 5-8** [smartphone]). Within the holding cage dashboard, new holding cages can be added using the “+” icon. The cage and mouse characteristics can be added based on the features mentioned previously under the database architecture. Individual cages can be viewed (“eye” icon), edited (“pencil/writing” icon) (**Supplemental video 9**), or deleted (“trash can” icon) (**Supplemental video 10**). Under the view menu, cage and mouse details can be viewed, along with a helpful “Sticky note” feature to note any special remarks regarding the cage or mice. Under the edit menu, individual cage or mouse characteristics or features can be edited and saved. We also included separate notes specific to individual mice which can be helpful when mice are earmarked for experiments or to note any odd behaviors or health issues for individual mice that may need follow-up. The “Maintenance logs” feature captures quick day-to-day entry logs for cage changes and other routine maintenance tasks (**Supplemental videos 11-12**). This feature can be used for multiple cages on the same page. An upload feature can be used to upload any files pertinent to the cage or individual mice such as genotyping gel image data (**Supplemental videos 13-14**). The breeding cage menu also has similar buttons within it with distinct editability for features that are different from holding cages such as litter and breeder mouse details (**Supplemental videos 15-16**).

A major shortcoming in recent open-source mouse colony management tools is the ability to quickly update cage records using mobile devices. To address this gap, we have included a feature that conveniently generates cage cards on ready-to-print templates (**Supplemental Figure 2a, Supplemental videos 17-18**). Each cage card template has a QR code that links it to its MyVivarium record (**Supplemental video 19**). Our print feature allows cage cards to be readily printed using a standard laser or deskjet printer on commercially available card stock with 3" x 5" perforations (4 per page) that can then easily fit in standard cage card holders (**Supplemental Figure 2b**).

Since vivarium work is closely linked to planning and scheduling experiments, we designed a “Tasks” feature to enable further integration of colony management into research projects. Individual tasks can be added using the “Manage tasks” menu under settings (**Supplemental videos 20-21**). We used an intuitive design to assign tasks to specific researchers and link these tasks to particular cages, with the option of adding due dates. A significant improvement compared to recent open-source solutions is the ability in MyVivarium to send automated email reminders related to tasks. Email reminders can conveniently be sent at user-defined frequencies. Once the user is logged in to their lab’s MyVivarium site, tasks appear on the homepage in a form similar to a Kanban board which can significantly enhance efficiency and productivity in project management [14].

While *a priori* design principles are important in determining the function of any given technology, design is critically linked to user experience and on-the-ground utilization of said technology [15,16]. In order to measure actual user engagement with a MyVivarium web application, we analyzed Google Analytics data collected over a period of four months (September 01, 2024 – December 31, 2024) from users of the Sathyanesan Lab MyVivarium (**Figure 3**). Our analysis of user metrics and engagement across specific pages within the application indicates significant user activity over this period (**Figure 3a-c**). In all three metrics measured, including number of active users (**Figure 3a**), views per user (**Figure 3b**), and user engagement in minutes (**Figure 3c**), users showed the highest levels of activity and engagement with holding cage-related pages, which is expected since most cages within our vivarium are holding cages and are most often used in experimental tasks. In order to assess the mode of user engagement, we analyzed MyVivarium usage by operating system (**Figure 3d**), device category (**Figure 3e**), and browser (**Figure 3f**). While certain categories were higher than others, we noticed that MyVivarium users had accessed the application from a diverse set of operating systems (Windows: 51.8%, Macintosh: 22%, Android: 13.1%, iOS: 10.1%, and Linux: 3%), device categories (desktop: 76.8%, mobile: 23.2%), and browsers (Chrome: 80.4%, Safari: 13.7%, Edge: 3%, Firefox 2.4%, Opera: 0.6%). Thus, MyVivarium works well as a pan-device, pan-OS, and pan-browser application.

**Figure 3.**
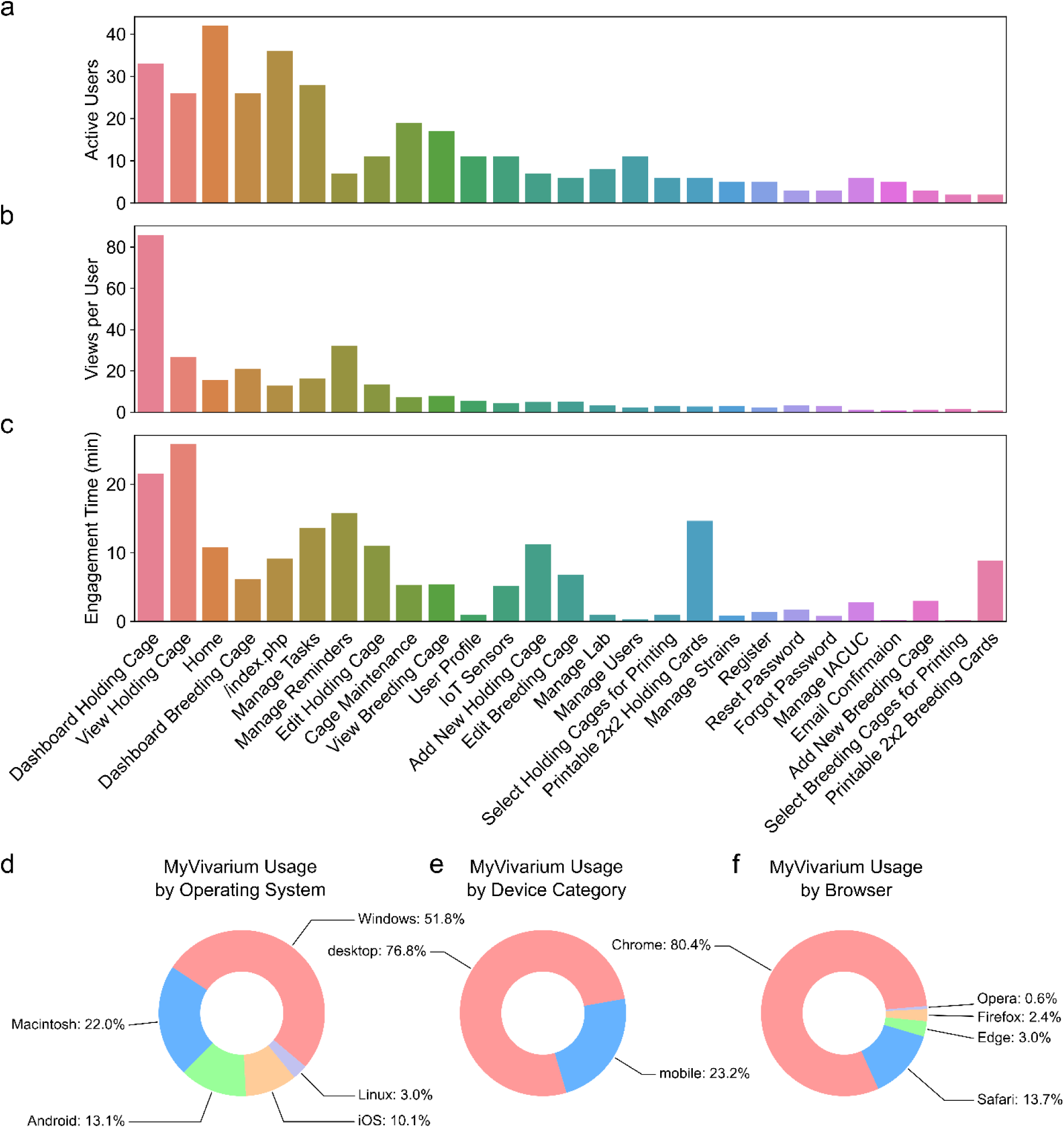
MyVivarium web application usage metrics for a single lab. **a.** Number of active users accessing the Sathyanesan Lab MyVivarium across different pages. **b.** Views per active user across page titles. **c.** Engagement time (in minutes) for a specific page. In panels **a-c,** note the highest metrics for holding-cage-related tasks since these are cages in higher number and are frequently used in experiments. Task management is also used significantly by users. **d.** MyVivarium usage percentage distribution by operating system, **e.** Device category, and f. Browser. In panels **d-f,** note the diversity in distribution across different categories, indicating pan-OS, pan-device, and pan-browser usability.

Since we included tools within MyVivarium to monitor the efficiency of vivarium management, we wanted to determine if and how these tools were being used. The “Manage Tasks” page was the fourth highest in terms of number of active users (**Figure 3a**) and fifth highest in terms of user engagement (**Figure 3c**). Among the many tasks for which users may employ the task management tool, weaning litters and conducting developmental experiments need careful adherence to timelines. We downloaded the database records as a CSV file using the “Export CSV” tool and analyzed management of these tasks (**Figure 4**). Out of the 63 tasks in the CSV, a total of 16 completed tasks fell under the weaning (8 tasks) and developmental experiment (8 tasks) categories. We used the task completion date as our proxy of when the experiments were actually completed. Our analysis of the weaning timelines showed that while these task records were created at different times following the birth of a litter, 100% of these tasks were completed by the target date (**Figure 4a**; day 0 on the relative x-axis or postnatal day 21 – P21). For the developmental experiment timelines, we found that four of the tasks (1, 2, 7, and 8) were completed earlier by 1 – 2 days (**Figure 4b**). Thus, our task management tool enabled a significant level of adherence to the target timeline, enabling compliance with vivarium rules, animal wellbeing, and scientific productivity.

**Figure 4.**
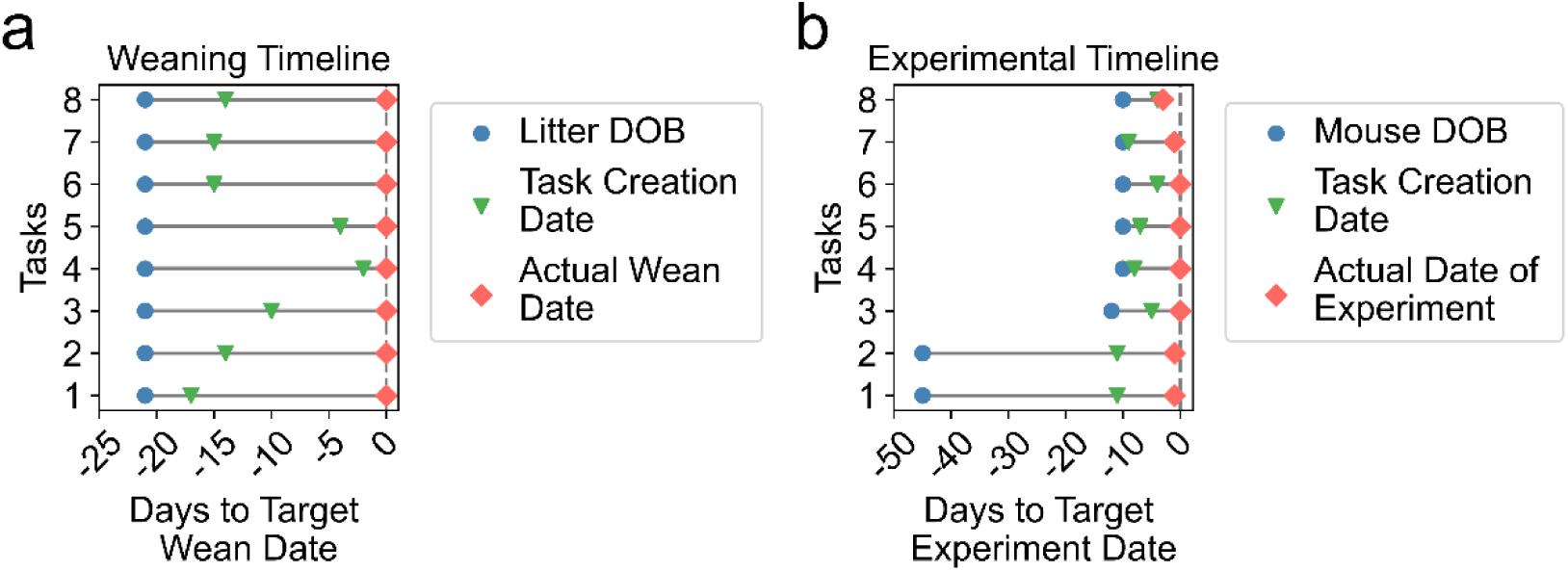
Task management ensures adherence to weaning and experimental timelines. **a.** Eight weaning timelines for which tasks were created for efficient management at differ ent times (green upside-down triangle; Task Creation Date) between litter date of birth (blue circle, Litter DOB) and actual wean date (red diamond; Actual Wean Date). **b.** Eight experimental timelines for which tasks were created (green upside-down triangle; Task Creation Date) also at different dates across the timeline. Dashed line in panels a and b denote the target day (day 0) for weaning (in **a**) or a timed experiment (in **b**). Note that in **a**, as inferred from the task completion date all litters are weaned on the target wean date. In **b**, all experimental tasks are completed by target date.

Recently, the relationship between ambient housing conditions of rodents used in research studies and potential confounding effects on metabolism and physiology has attracted much attention [17–19]. It is important to monitor ambient environmental factors since fluctuations in these factors can inadvertently alter animal health and behavioral responses, potentially skewing data, which would be of particular relevance to studies in neuroscience, endocrinology, physiology, and metabolism research. However, no colony management solution so far has the capability for ambient sensing of environmental factors in the vivarium. We designed and assembled a standalone open-source IoT-based multi-sensor system that is capable of monitoring temperature, humidity, illuminance, and vivarium worker activity, and feed this data directly into MyVivarium. We used inexpensive but accurate Inter-Integrated Circuit (I^2^C) sensors to enable the monitoring of ambient factors (**Figure 5a-b**) that can have an effect on colony wellbeing, metabolism, and physiology.

**Figure 5.**
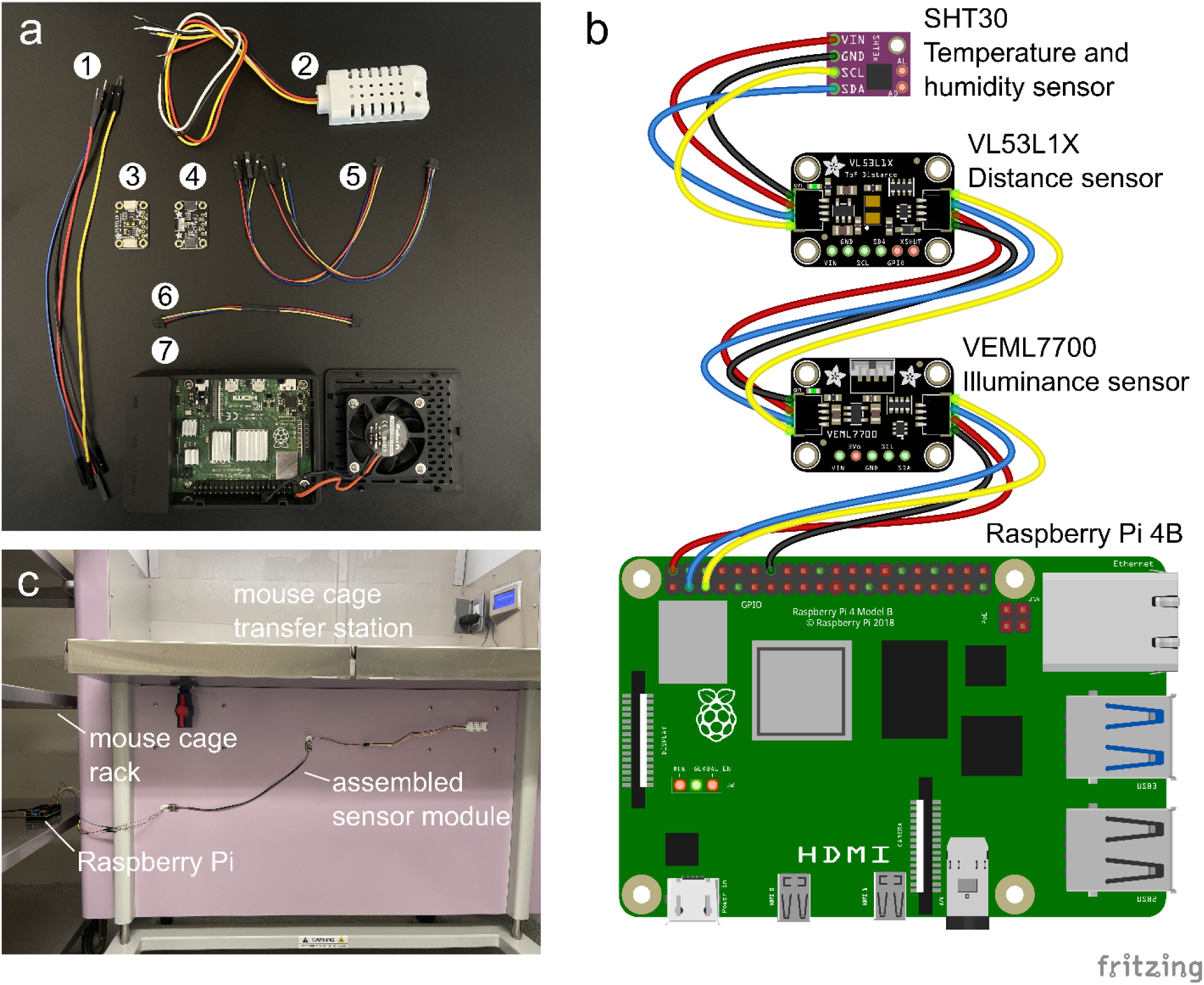
Design, assembly, and deployment of MyVivarium Pi-IoT sensor system. **a.** Components of Pi-IoT sensor system include 1: Jumper wires (to extend reach) 2: SHT30 Temperature and humidity sensor with plastic shell casing 3: VL53L1X distance or time-of-flight sensor which can measure distance to a person (vivarium-worker) 4: VEML7700 illuminance sensor which can detect lights ON and OFF 5 and 6: STEMMA-QT connectors (for Raspberry Pi GPIO pins to sensors and between sensors) 7: Raspberry Pi 4B (2 GB) in a case with a cooling fan. **b.** Fritzing sketch of serial connections between I^2^C sensors and Raspberry Pi GPIO pins based on a validated sensor module build. Wire color designations are as follows – red: power (+ve), black: ground (-ve), yellow: SCL (serial clock), blue: SDA (serial data). Note that in a, the color of SDA (data) is white. **c.** Assembled sensor module deployed in our vivarium room. Sensors have been affixed to the lower part of the mouse cage transfer station. The Raspberry Pi has been placed in a corner of a mouse cage rack.

Harnessing the ability of I^2^C sensors (targets) with unique addresses to communicate with a controller board synchronously, we serially connected a SHT30 (temperature and humidity), VL53L1X (proximity), and a VEML7700 (illuminance) sensor, each equipped with its own inbuilt pull-up resistor, to a Raspberry Pi 4B. We deployed a total of three sensor module sets (two in each of the vivarium rooms assigned to AS, and one in the vivarium room assigned to PP). In all three deployments, we placed the assembled sensor modules in the vivarium rooms in a configuration similar to that shown in **Figure 5c**. We used custom Python scripts to record ambient factors from data transmitted to the Pi, including temperature (°C), relative humidity (%), illuminance (lux), and vivarium worker-to-sensor proximity (cm) which is a proxy for vivarium worker presence or activity. We then used docker containers in IOTstack [20] to log sensor data to an InfluxDB database and visualize sensor datastreams using Grafana. We registered a low-cost domain name and used a Cloudflare tunnel to expose the local Grafana application to be accessible through this domain. We embedded the Grafana dashboard into MyVivarium using the inline frame element (iframe).

Using our RPi-IoT system, we have been able to record realtime ambient factor data continuously for months (**Figure 6**). In our Grafana dashboard we marked the maximum (“max”) and minimum (“min”) temperature (18-26°C) and relative humidity (30-70%) as recommended by *The Guide*, an authoritative publication issued by the office of laboratory animal welfare (OLAW), which is a part of the National Institutes of Health (NIH) [21], and the Jackson Laboratory, a non-profit mouse repository [22] for easy identification and reporting of alterations in these conditions. In general, temperature stayed within the recommended window (**Figure 6a**: data from four months, **Figure 6b**: data denoted in **Figure 6a** with arrows and grey bars from May 2^nd^ 2024), however, on few occasions across four months, relative humidity had short transient increases above the recommended maximum of 70% (**Figure 6c**). In general, increases in humidity seemed to follow the trend of relative humidity in Dayton, OH (**Figure 6c**, grey plot), with prolonged increases in external environmental humidity preceding transient spikes in vivarium rooms 1 (**Figure 6c**, **blue**) and 2 (**Figure 6c**, **orange**). Our illuminance sensor placements were able to effectively capture lights ON and OFF in rooms 1 and 2 (**Figure 6e-f**). Our proximity sensors were also able to detect activity in both rooms as inverted spikes (**Figure 6g-h**).

**Figure 6.**
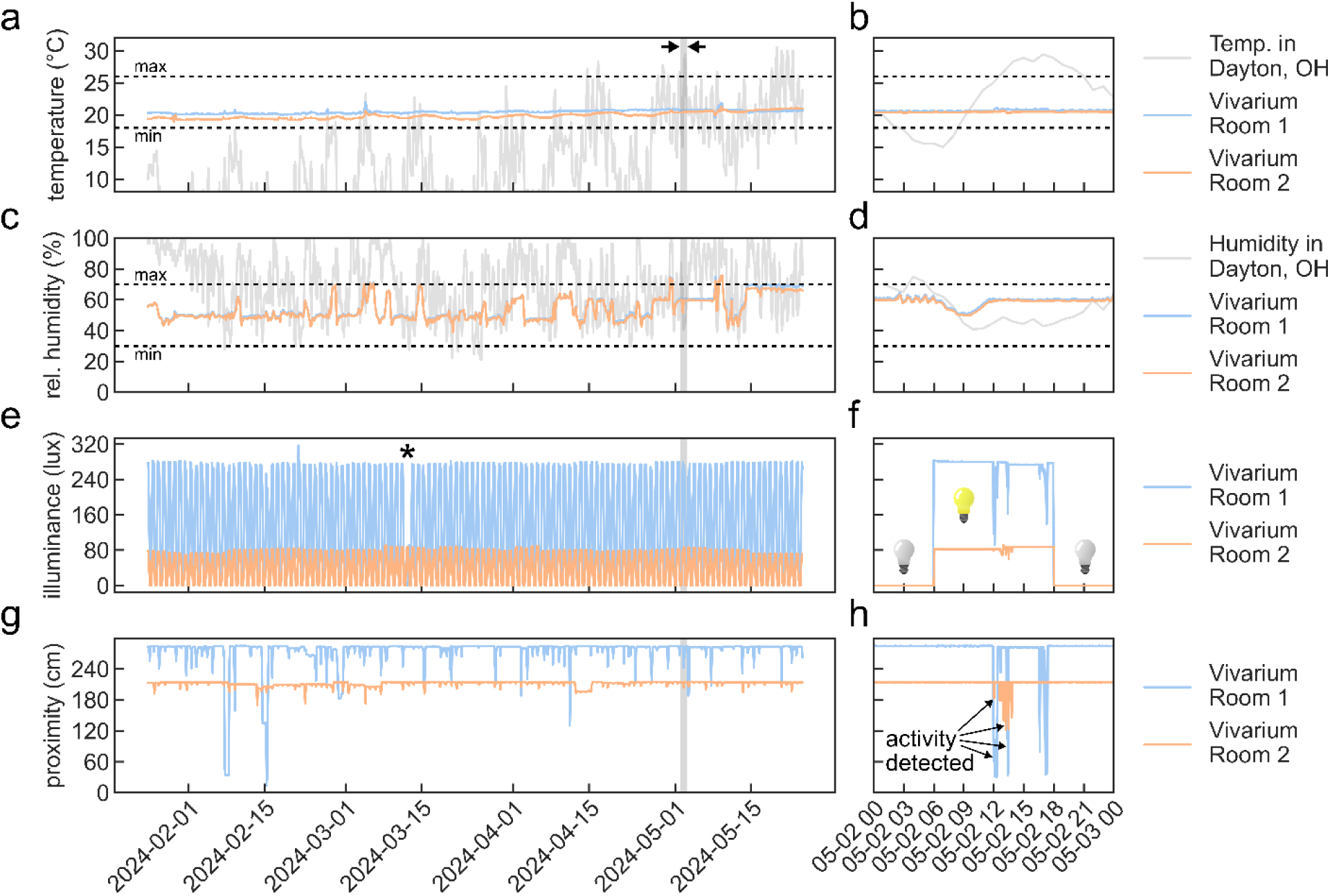
Continuous realtime ambient factor recording over a period of months using MyVivarium RPi-IoT sensor system. **a.** Temperature (°C) recorded over four months in vivarium rooms 1 (blue) and 2 (orange). Temperature in Dayton, OH is plotted as reference (grey). Dotted lines represent minimum (“min” = 18 °C) and maximum (“max” = 26 °C) recommended temperatures. Time-window (grey shaded rectangle) between arrows (12 midnight May 2^nd^, 2024 – 12 midnight May 3^rd^, 2024) is shown in greater detail in **b. c.** Relative humidity (%) recorded over four months. Humidity in Dayton, OH is plotted as reference (grey). Dotted lines represent minimum (“min” = 30%) and maximum (“max” = 70%). Time-window (grey shaded rectangle) is shown in greater detail in **d**. **e.** Illuminance sensor values recorded over months. Asterisk indicates an abnormal event detected by the sensor due to an erroneous manual override of the automated light cycle **f.** Sensor value change over the course of 24 hours shows vivarium lights OFF (light bulb turned off) and lights ON (light bulb turned on) in rooms 1 and 2 set to a 6 AM – 6 PM light cycle. **g.** Proximity sensor values for rooms 1 and 2 recorded for months **h.** Vivarium worker presence or activity is detected as negative deflections. Minimum and maximum temperature and humidity thresholds are based on suggested recommendations by *The Guide* (Office of Laboratory Animal Welfare, National Institutes of Health) and the Jackson Laboratory [21,22].

Using our system, we were able to monitor changes in ambient temperature, humidity, and illuminance (**Supplemental videos 22-23**). In response to increased humidity that was observed for a specific day (**Figure 7a**), we reported this to all users of our MyVivarium as a general message using “Sticky Notes” (**Figure 7b**). While this is an easy way to notify users of any abnormal events, it requires users to actively monitor the dashboard values. Our illuminance sensor values on March 12^th^ 2024 indicated that the lights in room 1 did not come ON as usual (**Figure 6e**, **asterisk**) due to an erroneous manual override of the automated light cycle. This error had not been identified and rectified till later that day. In order to reduce the likelihood of such events that could significantly alter the mood and wellbeing of mice, we set up an automated alerting system. We used the alerting system within Grafana to define thresholds for illuminance sensor values. We set two separate alert rules to fire corresponding to abnormal events within distinct phases of the light cycle i.e., lights ON > 15 minutes during the 6 PM – 6 AM window and lights OFF > 15 minutes during the 6 AM – 6 PM window. After confirming that the alert was working within Grafana, we created a dedicated Google Space to receive alerts via an integrated webhook. The webhook application program interface (API) is then used by the Grafana alerting system to send a predefined chat message with a link to the dashboard directly into the Google Space when the alert fires [23]. We tested our alerting system by blocking light entry to the illuminance sensor using black unreflective paper and were able to successfully receive alerts to the Google Space (**Figure *7*c-d**, **Supplemental video 24**). We have provided all code (https://github.com/myvivarium/RPi-IoT) and documentation (https://github.com/myvivarium/RPi-IoT/wiki) needed to set up our RPi-IoT system. An example snapshot of our Grafana interactive dashboard can be found here: https://snapshots.raintank.io/dashboard/snapshot/BS9oMWCz8rpT2H3xoGVoHyDHSobyJrrW

**Figure 7.**
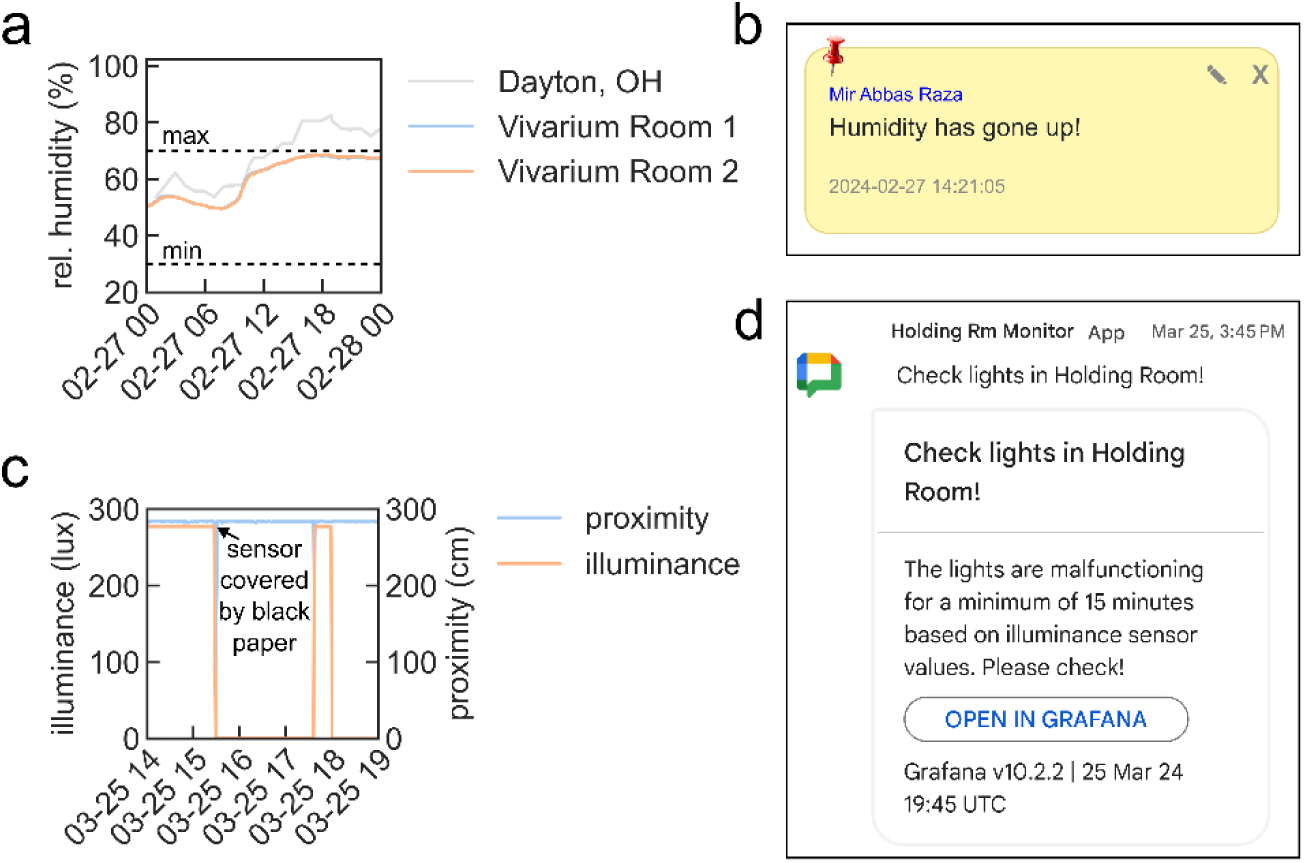
User notification of sensor readings and automated alerts. **a.** Increased humidity in vivarium rooms 1 (blue) and 2 (orange) on February 27^th^, 2024. **b.** General user notification using the “Sticky notes” on MyVivarium. **c.** Abnormal illuminance event and automated alert testing (arrow indicating when black unreflective paper was used to block the illuminance sensor from light exposure) shows reduced illuminance (orange) at 3:30 PM on March 25^th^ **d.** Alert message triggered by Grafana and relayed via a Google chat webhook app to a dedicated Google Space.

For admin-related tasks, we used a tier-based approach to control management access to MyVivarium. Typically, when setting up MyVivarium for the first time, a principal investigator (PI), or lab director/manager would designate themselves as an administrator (admin-tier, full access) and delegate one or a few key personnel as admins (admin-tier, full access) (**Supplemental video 25**). This enables distributive administration to increase efficiency of approving authorized users, determining access levels, and deleting unused or old user profiles. In addition to the admin-tier which has full access privileges, we have included a standard-tier user type with limited access to modifying and editing cage, mouse, and litter records. This enables greater control over which vivarium users have record-modifying privileges in order to reduce the likelihood of erroneous record modification. Admins can also perform strain management wherein new strains can be added to MyVivarium (**Supplemental videos 26-27**). This is advantageous for multiple reasons including standardization of mouse strain designation in cage and mouse records to prevent non-standard strain nomenclature by users, and easy linking of mouse strains to repositories such as The Jackson Laboratory to enable streamlining of mouse purchases or – in the case of custom generated lines – to enable sharing mouse lines with collaborators. We have also included and would like to emphasize the use of the *Research Resource Identifier* (RRID) for mouse strains to enable precise and standardized record-keeping [24]. Once strains have been added, users can easily search for these strains in the search bar when adding and filling out a cage record. Standardizing mouse strains in this manner helps researchers in both day-to-day research activities and in the sharing of results with the broader community of practice, especially for cases when a single mouse model can be represented by multiple strains in mouse repositories. In practice, since mouse strains and mouse models are often referred to by common names and not their lengthy official names or strain numbers listed in repositories, we have included common names to link particular strains to popular usage. As an example, in our mouse colony we house *Ts65Dn* mice which are an established Down syndrome mouse model [25]. Ts65Dn mice are generally available as two separate strains through Jackson Laboratory – strain #005252 [26,27] and strain #001924 [28,29]. These strains have official names B6EiC3Sn.BLiA-Ts(17^16^)65Dn/DnJ and B6EiC3Sn a/A-Ts(17^16^)65Dn/J respectively. “Ts65Dn” is the common name of both these strains. Using the combination of the RRIDs, official, and common names in MyVivarium helps disambiguate these strains in colony management. Finally, in order to enable integration of IACUC-related tasks and compliance such as assigning mouse strains to approved IACUC protocols and providing IACUC protocol access to users, we included a functionality where PIs or designated supervisors with admin access can perform these tasks in a streamlined way. PIs can easily assign mouse strains and upload IACUC protocols to their MyVivarium web applications (**Supplemental video 28**). We also included a “Export CSV” functionality to export all database tables in CSV format for download (**Supplemental video 29**), which can be helpful to monitor usage of mice and enable greater compliance with the 3Rs [30] as well as for data analysis regarding mouse breeding efficiency and correlation to ambient environmental factors.

Finally, we checked for potential objective performance issues on the client and server end. To check for client-side performance issues, we used GTmetrix (gtmetrix.com) which is a robust and widely-used service to test and measure web application performance (**Supplemental data file 1**; Test server location: Vancouver, Canada). The Sathyanesan Lab MyVivarium web application scored a grade of A with a 100% score on performance and 95% score on structure. This representative report validates high performance of our web application on the client side. On the server-side, we obtained key metrics from our cloud server platform (Cloudways) including unused (idle) CPU and unused (free) memory for one month (December 8^th^, 2024 – January 8^th^, 2025) (**Supplemental figure 3**). Despite our server running three independent MyVivarium web applications, both idle CPU (**Supplemental figure 3a**) and free memory (**Supplemental figure 3b**) was well within the recommended range (> 20% for idle CPU and > 100 MB free memory respectively). Thus, we validated both client-side and server-side performance for a MyVivarium web application with high performance scores.

## DISCUSSION

Mouse colony management is an integral part of biomedical and preclinical research [31]. There is a positive trend towards technology solutions for laboratory data management. The increasing popularity of electronic laboratory notebooks is a good example [32]. However, animal colony management has lagged behind. Two major reasons why technological solutions are not applied to certain laboratory tasks include cost and difficulty of implementation. We designed MyVivarium as an easy and low-cost solution to start and maintain a cloud-based cross-device colony management application. We have tested and validated the use of MyVivarium in three research labs. These include the Sathyanesan Lab, the Pitychoutis Lab, and the Goldstein Lab. The Sathyanesan and the Pitychoutis Labs are located in the University of Dayton (Dayton, OH), and the Goldstein lab is located in Augusta University (Augusta, GA). In our public repository we have provided detailed documentation for getting a MyVivarium web application up-and-running in a plug-and-play manner with minimal coding requirements. Our open-source RPi-IoT system can be used to stream realtime ambient environmental data from the vivarium rooms into a MyVivarium web application. In comparison to other tools (**Table 1**), MyVivarium is a significant step forward in enabling efficient animal colony management that is also open-source.

**Table 1.**
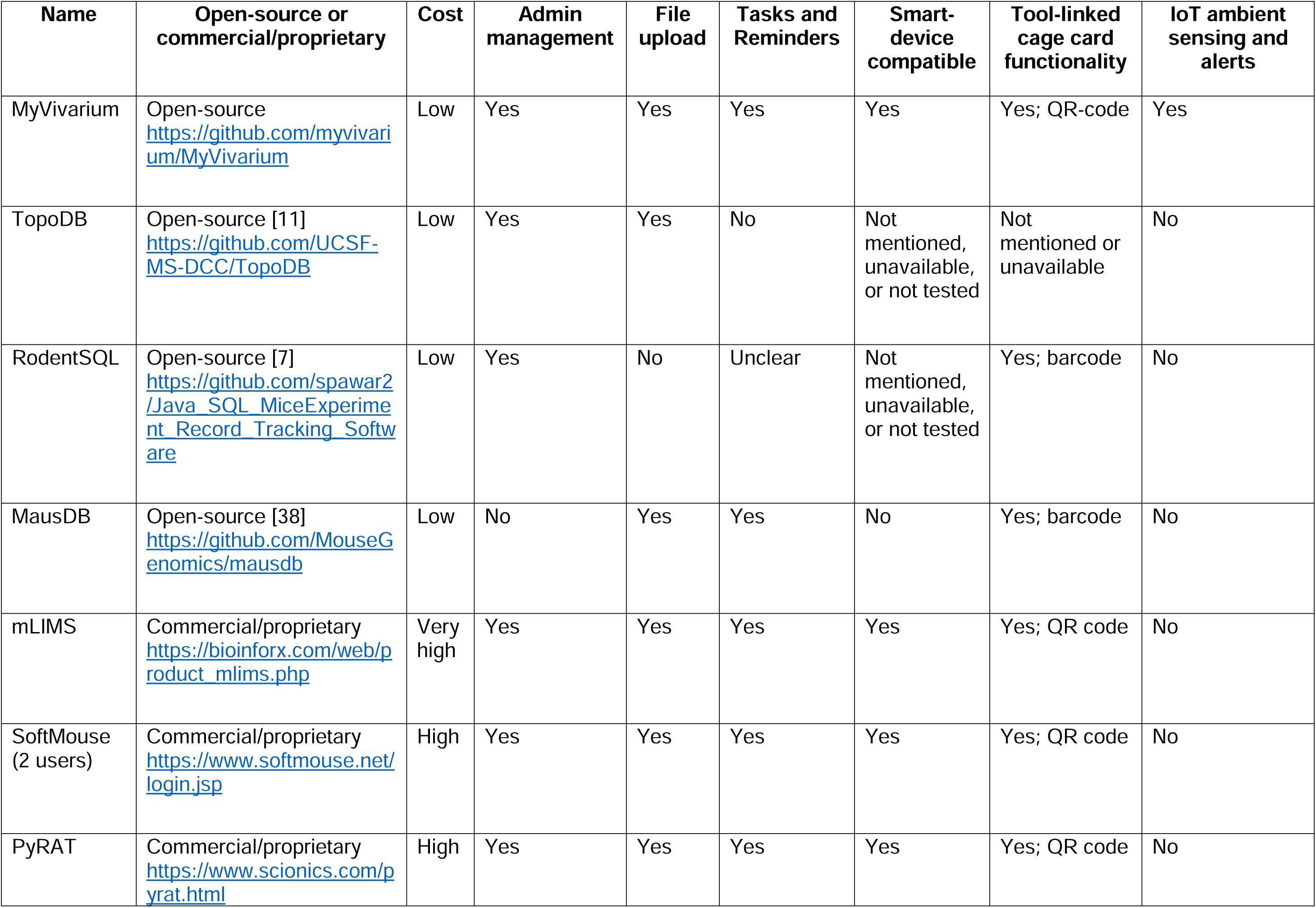
Comparison of MyVivarium to other available open-source and commercial colony management tools.

Our initial production deployment of MyVivarium for the mouse colony designated to AS at the University of Dayton was in October 2023, with a preliminary pilot version launched a few months prior in August 2023. We launched a second MyVivarium web application on the same server for the mouse colony designated to PP (also at the University of Dayton) in December 2023. A third MyVivarium web application was launched at Augusta University for the mouse colony designated to EG in November 2024. The two separate MyVivarium web app databases assigned to AS and PP at the University of Dayton have been tested with > 100 cage records across the two different labs with a combined total of > 250 mice belonging to 12 transgenic and three background lines. For the AS and PP labs, 15 users in total have access with 6 users having admin access. The MyVivarium web app database assigned to EG has been tested with 20 cage records with a total of 64 mice belonging to 5 transgenic lines. Across both institutions where MyVivarium has been tested, we have had seamless access to without any interruptions since deployment.

We employed the widely used and open-source LAMP stack (<u>L</u>inux as web server; <u>A</u>pache as database server; <u>M</u>ySQL as database; <u>P</u>HP as programming language) [33] to build MyVivarium. Even though the LAMP stack traces its origins to the early days of the modern web [34], its flexibility and open-source nature has made it a foundational set of web development tools still popular today. The LAMP stack is slower than the MEAN stack (MongoDB as database program; Express.js as web application framework back end; Angular as client-side application framework; Node.js as application runtime) in an “address book” system [35]. However, LAMP is superior to MEAN in handling relational data which is found in an animal colony management scenario [35]. While we have used widely used web components that have had relatively early origins, concerns about outdated forms of technology and digital objects are not entirely unfounded [36,37]. Many open-source software tools could potentially face issues due to a lack of active development. This is evident in the case of colony management software. To our knowledge, of the four significant publicly available open-source colony management tools including MausDB [38], JCMS [39], RodentSQL [7], and TopoDB [11], two are effectively unusable due to unavailability of critical installation files (MausDB), and potential incompatibility and obsolescence issues (JCMS) [40]. Additionally, while the available MausDB public repository has detailed documentation regarding installation, JCMS lacks any installation documentation with no active forks in development by other users. Given these realities, the fact remains that open-source remains an effective strategy in combating digital obsolescence [41].

To make MyVivarium as accessible as possible, we prioritized cost-savings in both web components as well as the hardware and software components needed for the RPi-IoT system. Our initial server costs were significantly reduced due to discounts from our hosting provider ($0 for 6 months). Similar to our setup, if a “founder lab” sets up a server using a cloud hosting provider, this single server can then be used to launch multiple MyVivarium applications for multiple other labs that want to use MyVivarium. This would be an additional cost-saving measure, reducing the server cost further depending on the number of labs that join the server. Importantly, we designed MyVivarium such that each lab would have their own secure instance without their data being accessed by the lab that initially set up the server. When comparing costs of similar systems to monitor IoT in vivarium-related contexts (**Table 2**), we found that our RPi-IoT system was significantly cheaper than the next highest priced system – the MR1 sensor [42]. It must be mentioned however, that the comparison between our system and the MR1 sensor is not strictly one-to-one since the MR1 is placed within a cage and hence captures ambient data within a cage and captures mouse activity instead of vivarium worker activity. Although our system can readily be modified to this end (the presence sensor in our system is similar in function to what is likely a PIR sensor in the MR1), this is not exactly our aim at this time. In addition, the free version of the cloud interface provided by Pallidus Sensing LLC only allows users 30 prior days of retrieval [43]. Our dashboard design using the Grafana OS version allows for unlimited retrieval of data.

**Table 2.** Comparison of MyVivarium RPi-IoT ambient sensing tool compared to other tools.

| Name | Open-source or commercial/proprietary | Cost of components (in USD) |  |
| --- | --- | --- | --- |
|  |  | Software-related | Hardware-related |
| MyVivarium RPi-IoT system | Open-source<br>https://github.com/myvivarium/RPi-IoT | Grafana Open source: \$0;<br>Domain name registration: varies, but as low as \$6 per year | \$110.08 |
| Pallidus MR1 sensor and Pallidus cloud software (Pallidus Sensing LLC) | Commercial/proprietary<br>https://pallidus.io/products/mr1-sensor | Pallidus Free: \$0 (only access to last 30 days of data); Pallidus professional cloud: \$2,990.00 | \$399.00 per sensor, however can only be added to an existing installation; Pallidus starter kit (8 sensors): \$4,499.00 (includes only one year subscription to cloud) |
| REM Module and Guardian software (excludes software maintenance, training, and installation) (Techniplast, USA) | Commercial/proprietary<br>https://digitalcage-tecniplast.com/en/products/rem.html | > \$800.00 license fee | > \$2000.00 |

While we consider MyVivarium to be a significant advance in comprehensive open-source colony management, we would like to note some limitations of this tool. First, as is the case with all web applications that are hosted on the cloud, MyVivarium web applications can be potentially subject to malicious action by unapproved actors. Cloud security in a MyVivarium web application that is enhanced by Cloudflare, while significantly reducing the chances of attack, does not completely eliminate all likelihood of malicious entry. As best practice related to any web-related application, we provide three suggestions including the prioritization of database security by admins, obtaining regular backups of the database stored in a secure location, and strong password management by all users. While we built MyVivarium as being primarily remote-accessible, users with primary concerns over security can also choose to host a MyVivarium web application running on a local server. Second, a MyVivarium cloud-hosted web application is also subject to internet outages which are becoming more common according to recent analysis by Cloudflare [44]. Third, we have not included any version tracking of cage records triggered by user-defined changes. Since MyVivarium is under active development, we plan to incorporate this feature in a future release. Finally, the total number of cage records that we have included in our specific MyVivarium applications is lower compared to other tested solutions [11] emphasizing the need for further testing and performance monitoring as cage records increase. However, given the current seamless performance of MyVivarium in our hands, we do not anticipate significant performance deficits for other labs with hundreds of cages.

## CONCLUSION

MyVivarium is a low-cost, effective, and rapidly deployable cloud-based solution for research mouse colony management. Cloud-based access and the integration of open-source realtime ambient sensing sets MyVivarium apart as a major step forward in digital colony management. We envision MyVivarium to be of significant benefit to investigators who would like to make their colony management more efficient without expending much time or effort by themselves or their personnel. Future development includes the incorporation of MyVivarium into a larger ecosystem of lab management. This would enable the tracking of budgets, purchasing, and chemical inventory in addition to colony management. This integration into an easy-to-deploy open-source cloud-based electronic laboratory notebook currently in development. It would also be worthwhile to explore how sectors not directly related to vivarium work such as precision livestock farming (PLF) can contribute to improving the use of IoT in the vivarium [45,46]. Finally, we also plan to develop and incorporate tools for big data analysis into MyVivarium to enable researchers to identify critical factors that affect colony breeding efficiency, health, and well-being.

## METHODS

### Web-server and database building strategy

We followed a structured approach to build the web server and database for MyVivarium, ensuring the system is robust and user-friendly. The strategy encompasses installing and configuring the web server, integrating PHP for dynamic content generation, and setting up a reliable database system. We used PHP for server-side scripting, leveraging its ease of integration with web servers and widespread use in web development. The web server utilized is Apache, known for its robustness and extensive documentation, ensuring reliable performance. Composer, a dependency manager for PHP, is employed to manage libraries such as PHPMailer for email functionality and Dotenv for environment variable management. Configuration settings, including database credentials and SMTP settings, are stored in a ’.env’ file, which the Dotenv library loads into the application for secure and flexible configuration. MySQL is the database management system used, and it was chosen for its reliability and performance. The database schema manages various aspects of vivarium operations, including user management, breeding cages, holding cages, tasks, notes, and more. Tables are created with appropriate relationships and constraints to ensure data integrity.

#### Web server setup

We selected Apache as our web server because of its widespread usage and strong support in the industry. Apache offers a stable and secure platform that can effectively meet the needs of MyVivarium. The setup process entails installing Apache and configuring it to host the MyVivarium application. It involves setting up virtual hosts to manage multiple web applications on a single server and configuring the server to utilize PHP for processing dynamic web pages.

#### PHP integration

PHP is integrated into the Apache web server to enable the server to process PHP scripts. PHP is a widely used scripting language, especially suited for web development. It allows us to create dynamic content and seamlessly interact with the MySQL database. Composer, a dependency manager for PHP, handles external libraries and dependencies, ensuring the application has all the necessary tools and libraries to function correctly.

#### Database configuration

We chose MySQL as the database management system because of its reliability and efficiency in handling large amounts of data. The database schema stores and manages different data types needed for MyVivarium, including user information, breeding and holding cage data, tasks, and notes.

#### Environment configuration

The configuration settings for the database and other essential components are stored in a ’.env’ file. This file includes environment-specific settings, such as database credentials and SMTP settings for email functionality. The application utilizes the Dotenv library to load these settings, providing a secure and flexible method for managing configuration data.

#### Security and compliance

We placed security as a high priority in the development of MyVivarium. Our system is configured to ensure secure data transmission and storage. Passwords are hashed before being stored in the database, and sensitive information is protected using state-of-the-art encryption. Our system is designed to comply with industry standards and regulations, ensuring user data is handled with the utmost care.

#### Deployment

To deploy the application, we set up the web server and database on a server running Ubuntu, a widespread and stable Linux distribution known for its security. The process involves installing Apache, PHP, and MySQL, configuring the server to host the MyVivarium application, and ensuring all services run smoothly. After setup, we tested the application to ensure it was fully functional and ready for use. A detailed guide on installing a LAMP (Linux, Apache, MySQL, PHP) stack on Ubuntu can help new users set up their own version of MyVivarium [47]. We used Cloudflare as the DNS provider, adding another layer of security to our MyVivarium application. To implement this, we moved our domain from our previous registrar to Cloudflare by updating the domain’s nameservers to Cloudflare’s nameservers. This process involves logging into the domain registrar’s account, locating the DNS settings, and replacing the existing nameservers with those provided by Cloudflare. After the domain was successfully transferred, we subscribed to Cloudflare’s free plan, which includes essential security features such as DDoS protection and basic firewall rules. These measures protect the application from malicious attacks and ensure high availability. We configured Cloudflare to filter incoming traffic, block potential threats, and optimize content delivery for improved performance and security. This setup reasonably ensures that a MyVivarium web application remains resilient against cyber threats while maintaining seamless access for legitimate users. Note that this configuration needs to be done separately and is not part of the code in the GitHub repository. Finally, a complete list of all the files in MyVivarium have been included in our GitHub repository (https://github.com/myvivarium/MyVivarium).

### Cage-card print functionality and QR-code accessibility

In order to physically link cage-records to MyVivarium, we incorporated a cage-card print functionality. We printed cage-cards directly from a browser and onto card stock with 3" x 5" perforations (4 per page) (Amazon.com, WA; ASIN: B09BZST43T) using a standard LaserJet printer. We incorporated QR-codes into the cage-card design in order to increase efficiency in managing records using mobile devices. We have tested QR-code accessibility using android tablets, as well as android-based and apple-based smartphones. In all our testing and use, QR-code access has been seamless.

### RPi-IoT design and assembly

#### Hardware components

We used a Raspberry Pi based design for our IoT system. All sensors used were I^2^C based which made the use of multiple sensors with different addresses straightforward. For each of our sensor modules, we used a Raspberry Pi model 4B with 2 GB RAM (Adafruit Industries LLC, NY) with a 128 GB RAM running the 64-bit Raspberry Pi Bookworm OS (https://www.raspberrypi.com/software/). The Pi was cooled inside a case using both active cooling and passive cooling elements (Amazon.com, WA; B0963HHSWK). We used three I^2^C sensors including the SHT30 Temperature and Humidity Sensor (Adafruit LLC, NY), Adafruit VEML7700 Lux Sensor -I2C Light Sensor (Adafruit LLC, NY), and the Adafruit VL53L1X Time of Flight Distance Sensor for 30 to 4000mm (Adafruit LLC, NY). Connectors and wires were purchased from Adafruit. A detailed list of hardware components with vendor weblinks is provided in the bill of materials in our GitHub repository (https://github.com/myvivarium/RPi-IoT).

#### Data visualization and software components

Detailed steps regarding software installation steps can be found on our GitHub repository wiki (https://github.com/myvivarium/RPi-IoT/wiki). Briefly, we installed TigerVNC server (https://tigervnc.org/) on the Pi to enable remote sessions for easy management and data retrieval. We then configured the Pi to be able to detect I2C device interfacing. We also registered a free DuckDNS account to track changes in the IP address of the Pi in the event it restarted and a new IP address was assigned to the Pi. Next, we used IOTstack to run all necessary IoT-related applications on the Pi using a Docker stack [20]. Specifically, we built our stack to include DuckDNS, Grafana, InfluxDB, and Portainer. We then created a database: `multisensors` to store sensor data. We then installed all relevant libraries for the Pi to detect streaming data from the I2C sensors. We wrote a python script (`influxdb-multisensor.py`) to log sensor data to the `multisensors` database as well as store a redundant CSV per day of recording [48]. Intervals at which data can be logged to the InfluxDB database is user-defined, however, relevant for our purposes and also to balance sensor functionality, we used a time interval of 1 minute. After confirming that the database is actively being updated with sensor data input using Grafana, we installed a service file to ensure that the `influxdb-multisensor.py` script runs even if the Pi restarts. We then purchased a low-cost domain name using a domain name registration service (Cloudflare, San Francisco, CA; 1 year domain registration price = $6.65 USD), added this domain to Cloudflare using a free account, and used a Cloudflare tunnel [49,50] to securely expose the local Grafana application running on the Pi. Once the Cloudflare tunnel was active, we were able to access our sensor data directly from the domain. We then obtained the embed URL for individual panels of our dashboard and used these in MyVivarium under the Admin tasks section to directly view realtime sensor data. Finally, in order to enable alerting via Grafana and Google chat, we used a webhook app within Google Space and added this as a contact point in Grafana’s alerting system. We defined alert rules for abnormal illuminance sensor values for both light and dark cycles of the vivarium rooms. We set the evaluation period as 15 minutes with a pending period of 15 minutes, which meant that if the sensor was detecting abnormal light function for at least 30 minutes, then it will send a notification via the webhook app.

### Vivarium facility

The University of Dayton vivarium is a modern animal facility specifically designed for rodent (mice and rats) micro-isolator caging with each room that holds animal cages containing Allentown Phantom animal transfer stations. All rooms including those that hold animal cages as well as procedure rooms are independently controlled for air pressure (positive or negative), temperature, and timer-controlled lighting. Lighting in rooms include both white and red dimmable lights. Humidity is controlled centrally by the University of Dayton facilities management. The Augusta University vivarium is located in the building adjacent to the Goldstein Laboratory. The Division of Laboratory Animal Services (DLAS), accredited by the Association for Assessment and Accreditation of Laboratory Animal Care International (AAALAC International), is primarily responsible for the daily maintenance and veterinary care of the colony, and are available 24/7. All mice in the colony are housed in an Allentown 70-cage rack that are also monitored by an Ecoflo system, regulated by DLAS. Lighting is controlled by an Intermatic system on a 12-hour on/off schedule, while humidity and temperature are monitored by an Acurite meter, with adjustments made by DLAS staff as necessary. While the DLAS is responsible for day-to-day tasks such as cleaning cages, feeding/watering, and acknowledging and treating injuries according to the standards of the AAALAC International, members of the Goldstein Lab are tasked with the upkeep of weaning schedules.

## ACKNOWLEDGEMENTS

We would like to thank the Director of the University of Dayton Vivarium, Dr. Jeffrey Kavanaugh, and the University of Dayton vivarium veterinarian, Dr. Paul A. Stull, DVM, for helpful suggestions during the early phase of this study.

## CRediT authorship contribution statement

**Robinson Vidva:** Conceptualization, Methodology, Software, Visualization, Writing – review and editing, Project administration. **Mir Abbas Raza:** Conceptualization, Investigation, Data Curation, Methodology, Writing – review and editing, Project administration, Visualization. **Jaswanth Prabhakaran:** Conceptualization, Methodology, Software. **Ayesha Sheikh:** Data Curation, Investigation, Writing – review and editing. **Alaina Sharp:** Data Curation, Investigation, Writing – review and editing. **Hayden Ott:** Data Curation, Writing – review and editing. **Amelia Moore:** Data Curation. **Christopher Fleisher:** Data Curation. **Hailey Netherton:** Validation, Data Curation. **Evan Goldstein:** Project administration, Validation, Resources, Writing – review and editing. **Pothitos M. Pitychoutis:** Project administration, Resources, Writing – review and editing. **Tam V. Nguyen:** Conceptualization, Methodology, Writing – review and editing, Project administration, Supervision. **Aaron Sathyanesan:** Conceptualization, Methodology, Software, Formal analysis, Resources, Writing – review and editing, Writing – original draft, Visualization, Project administration, Supervision, Funding acquisition.

## DATA AND CODE AVAILABILITY

All code has been uploaded to public GitHub repositories: https://github.com/myvivarium/MyVivarium and https://github.com/myvivarium/RPi-IoT. Data included in this manuscript has been uploaded to the open science foundation (OSF) data repository and can be downloaded using this link: https://osf.io/7x8tc/?view_only=c3e4433a2ac64103a8f33c71ccd7504a

## Footnotes

1 Client-side refers to end-user devices, like laptops and smartphones. Server-side refers to the devices that manage a web application.

