## Supplemental Figure 1 for "MyVivarium: A cloud-based lab animal colony management application with realtime ambient sensing"

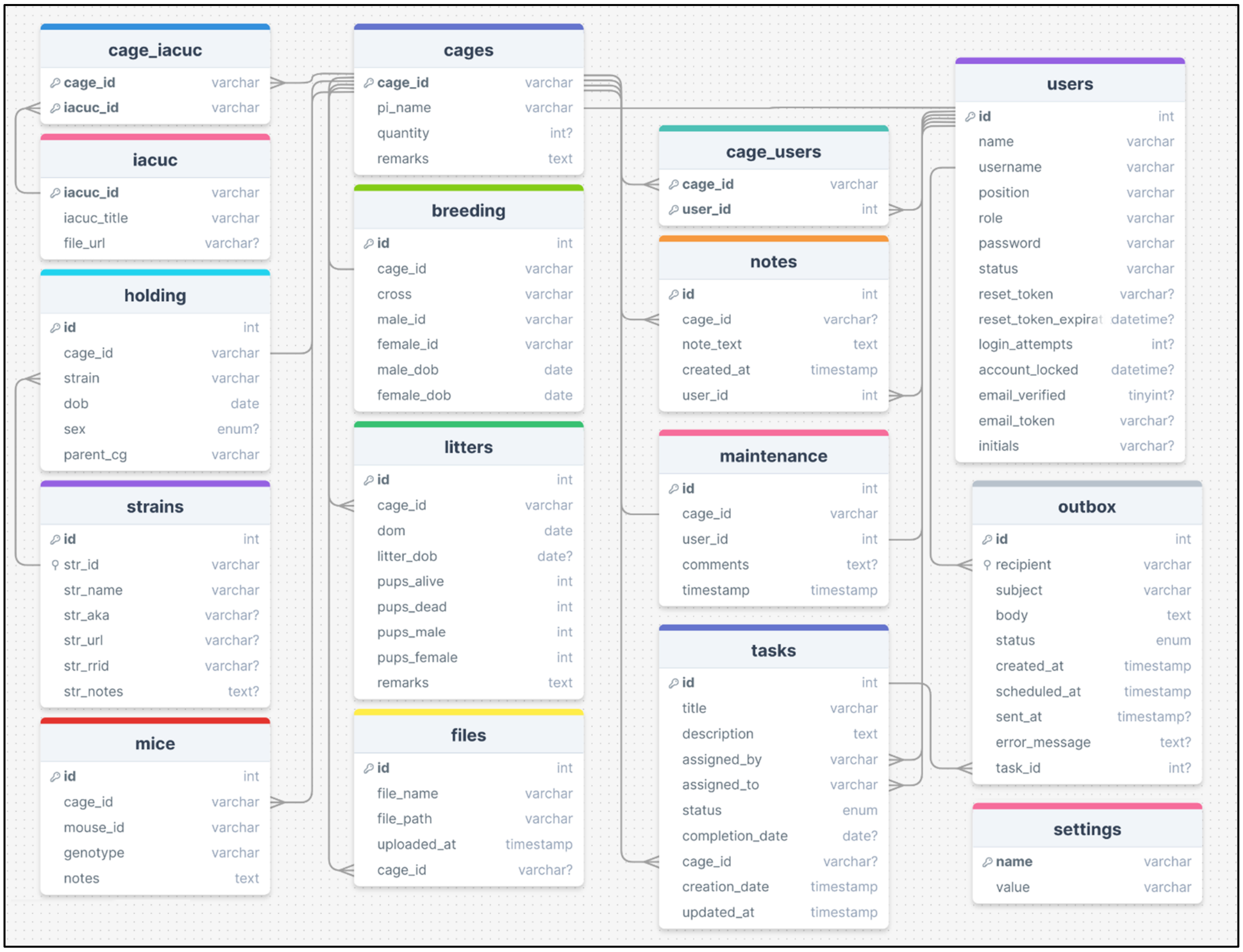


**Supplemental Figure 1.** The entity relationship diagram (ERD) for MyVivarium describes its database system for managing laboratory animal cages, mice, breeding, and related tasks. Key entities include `cages`, which store details about each cage, and `iacuc`, which provides information on associated protocols. The `cage_iacuc` entity connects cages to IACUC protocols, while `cage_users` links users to specific cages. `holding` tracks mouse information such as strain and sex, and `strains` contains detailed strain data. The `mice` entity records data on individual mice, including genotype. `breeding` records breeding activities, and `litters` track litter details. `files` manage documents associated with the cages, and `users` contain user information. The `notes` entity stores notes related to cages, `maintenance` logs maintenance activities, `tasks` manages task assignments, `outbox` tracks communication records, and `settings` stores system configurations. Fields marked with '`?`' indicate nullable fields, meaning they may not always contain data.
