## Supplemental Figure 2 for "MyVivarium: A cloud-based lab animal colony management application with realtime ambient sensing"

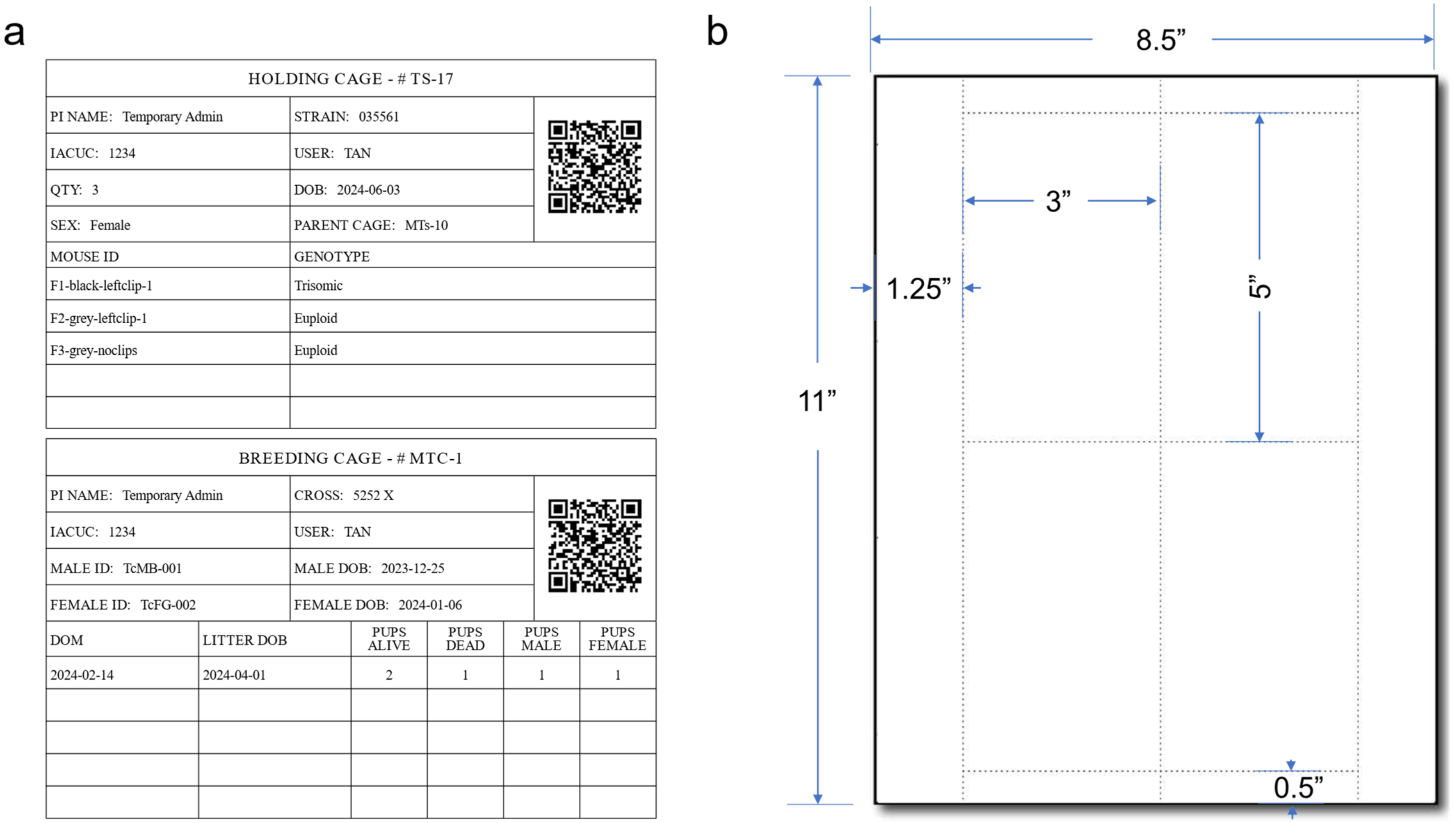


**Supplemental Figure 2. a.** An example of two cage cards printed using the print functionality in MyVivarium. Each cage card can be customized according to the needs of individual labs, however, by default, each card has a dimension of 3”×5”. Scanning the QR code links to the cage records in a demo MyVivarium web application which can be accessed using the demo credentials on the landing page **b.** Template for printing four cage cards onto pre-perforated card stock with default dimension.
