## Supplemental Figure 3 for "MyVivarium: A cloud-based lab animal colony management application with realtime ambient sensing"

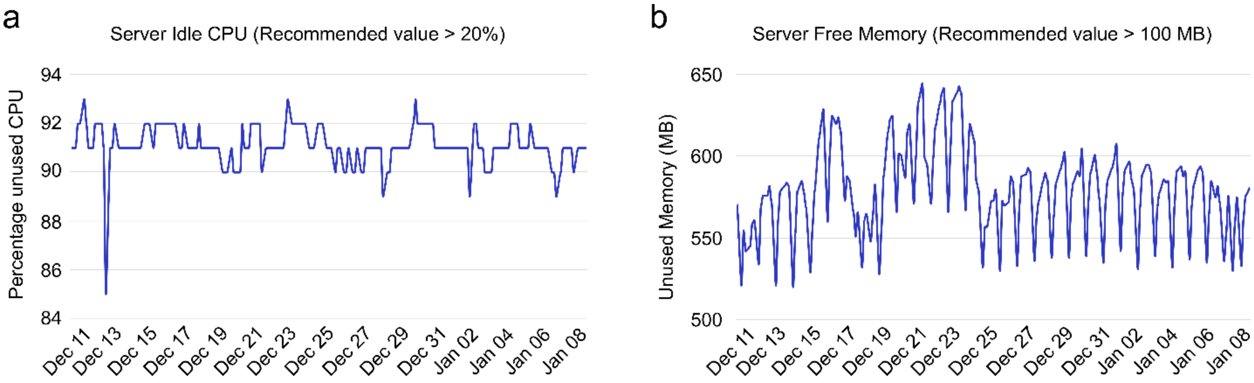


**Supplemental Figure 3. Server usage and performance metrics running three MyVivarium web applications. a.** Percentage of idle CPU (unused CPU) for our Cloudways server between December 8th 2024 – January 8th 2025. Note that the percentage of unused CPU is always above the recommended value of 20% **b.** Unused (free) server memory for the same time period. Note that the percentage of unused memory is always above the recommended value of 100 MB.
