## Supplemental data file 1 for "MyVivarium: A cloud-based lab animal colony management application with realtime ambient sensing"

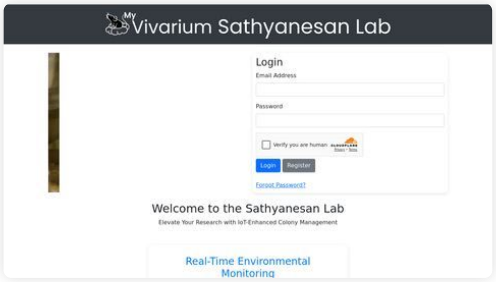


### Performance Report for:

#### https://myvivarium.online/

Report generated: Mon, Jan 6, 2025 11:49 AM -0800
Test Server Location: Vancouver, Canada
Using: Chrome 117.0.0.0, Lighthouse 11.0.0

| GTmetrix Grade | Performance | Structure | L. Contentful Paint | T. Blocking Time | C. Layout Shift |
|---|---|---|---|---|---|
| A | 100% | 95% | 442ms | 0ms | 0 |

#### Top Issues

| Priority | Issue | Result |
|---|---|---|
| Med | **Use explicit width and height on image elements** CLS | 1 image found |
| Low | **Allow back/forward cache restoration** | 2 failure reasons |
| Low | **Eliminate render-blocking resources** FCP LCP | Potential savings of 67ms |
| Low | **Avoid chaining critical requests** FCP LCP | 5 chains found |
| Low | **Avoid enormous network payloads** LCP | Total size was 698KB |

##### Focus on these audits first

These audits likely have the largest impact on your page performance.

Structure audits do not directly affect your Performance Score, but improving the audits seen here can help as a starting point for overall performance gains.

#### Page Details

2.6s
**Fully Loaded Time**

##### Total Page Size - 696KB

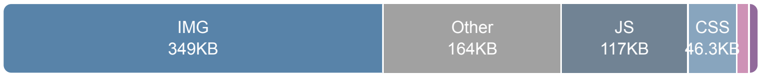


##### Total Page Requests - 35

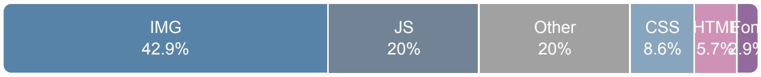


HTML JS CSS IMG Video Font Other

##### How does this affect me?

Modern web users have a short attention span and expect a fast and seamless website experience. Delivering that fast experience can result in more traffic, more conversions, and more happiness.

As if you didn't need more incentive, **Google use Page Speed and Page Experience (including Web Vitals) signals in their ranking algorithm.**

##### About GTmetrix

GTmetrix was developed as a tool for customers to easily test the performance of their webpages.

[Learn more about us.]

The waterfall chart displays the loading behaviour of your site in your selected browser. It can be used to discover simple issues such as 404's or more complex issues such as external resources blocking page rendering.

**Sathyanesan Lab**

| | | | | |
|---|---|---|---|---|
| / | 200 | myvivarium.online | 4.98KB | 159ms |
| **bootstrap.min.css** | 200 | maxcdn.bootstrapcd… | 25.0KB | 70ms |
| **bootstrap.min.css** | 200 | cdn.jsdelivr.net | 20.4KB | 18ms |
| **css2?family=Poppins:wg…** | 200 | fonts.googleapis.com | 999B | 44ms |
| **jquery.min.js** | 200 | ajax.googleapis.com | 30.3KB | 42ms |
| **bootstrap.min.js** | 200 | maxcdn.bootstrapcd… | 15.3KB | 83ms |
| **logo1.jpg** | 200 | myvivarium.online | 117KB | 158ms |
| **DSC_0536.webp** | 200 | myvivarium.online | 13.7KB | 177ms |
| **DSC_0537.webp** | 200 | myvivarium.online | 15.4KB | 316ms |
| **DSC_0539.webp** | 200 | myvivarium.online | 16.8KB | 185ms |
| **DSC_0540.webp** | 200 | myvivarium.online | 15.7KB | 191ms |
| **DSC_0560.webp** | 200 | myvivarium.online | 22.2KB | 201ms |
| **DSC_0562.webp** | 200 | myvivarium.online | 13.2KB | 206ms |
| **DSC_0586.webp** | 200 | myvivarium.online | 31.3KB | 207ms |
| **DSC_0593.webp** | 200 | myvivarium.online | 23.8KB | 206ms |
| **DSC_0607.webp** | 200 | myvivarium.online | 21.4KB | 206ms |
| **DSC_0623.webp** | 200 | myvivarium.online | 11.2KB | 205ms |
| **DSC_0658.webp** | 200 | myvivarium.online | 22.5KB | 206ms |
| **DSC_0665.webp** | 200 | myvivarium.online | 23.7KB | 207ms |
| **api.js** | 302 | challenges.cloudflar… | 0 | 152ms |
| **vcd15cbe7772f49c399c6a…** | 200 | static.cloudflareinsig… | 6.93KB | 138ms |
| **pxiEyp8kv8JHgFVrJJfecg…** | 200 | fonts.gstatic.com | 7.70KB | 50ms |
| **s.js?z=JTdCJTIyZXhlY3V…** | 200 | myvivarium.online | 4.28KB | 24ms |
| **api.js** | 200 | challenges.cloudflar… | 15.7KB | 36ms |
| **auto/** | 200 | challenges.cloudflar… | 7.39KB | 46ms |
| **1** | 200 | challenges.cloudflar… | 61B | 19ms |
| **v1?ray=8fde3b6c7a722d6…** | 200 | challenges.cloudflar… | 44.5KB | 21ms |
| **manifest.json** | 200 | myvivarium.online | 933B | 105ms |
| **POST rum?** | 204 | myvivarium.online | 0 | 13ms |
| **favicon.ico** | 200 | myvivarium.online | 8.19KB | 89ms |
| **android-chrome-192x192.…** | 200 | myvivarium.online | 20.5KB | 125ms |
| **POST ywh3HEjgEvz4GFfj…** | 200 | challenges.cloudflar… | 113KB | 97ms |
| **zvUR3mihxkyLL09** | 401 | challenges.cloudflar… | 1.66KB | 31ms |
| **C6fQjKlzdjvGWKr** | 200 | challenges.cloudflar… | 180B | 18ms |
| **POST ywh3HEjgEvz4GFfj…** | 200 | challenges.cloudflar… | 19.6KB | 75ms |

**35 Requests** **696KB (1.54MB Uncompressed)** **2.6s (Onload 633ms)**

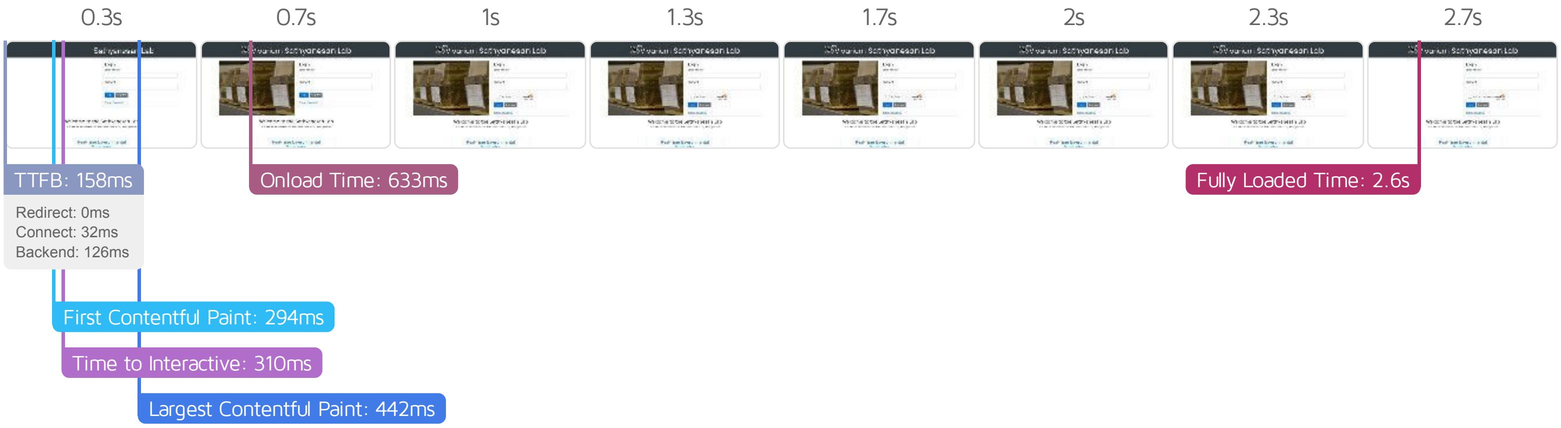


#### Performance Metrics

##### First Contentful Paint

How quickly content like text or images are painted onto your page. A good user experience is 0.9s or less.

Good - Nothing to do here

293ms

##### Time to Interactive

How long it takes for your page to become fully interactive. A good user experience is 2.5s or less.

Good - Nothing to do here

309ms

##### Speed Index

How quickly the contents of your page are visibly populated. A good user experience is 1.3s or less.

Good - Nothing to do here

374ms

##### Total Blocking Time

How much time is blocked by scripts during your page loading process. A good user experience is 150ms or less.

Good - Nothing to do here

0ms

##### Largest Contentful Paint

How long it takes for the largest element of content (i.e., a hero image) to be painted on your page. A good user experience is 1.2s or less.

Good - Nothing to do here

442ms

##### Cumulative Layout Shift

How much your page's layout shifts as it loads. A good user experience is a score of 0.1 or less.

Good - Nothing to do here

0

#### Browser Timings

| | | | | | |
|---|---|---|---|---|---|
| Redirect | 0ms | Connect | 32ms | Backend | 126ms |
| TTFB | 158ms | DOM Int. | 256ms | First Paint | 294ms |
| DOM Loaded | 310ms | Onload | 633ms | Fully Loaded | 2.6s |

| IMPACT | AUDIT | |
|---|---|---|
| Med | Use explicit width and height on image elements CLS | 1 image found |
| Low | Allow back/forward cache restoration | 2 failure reasons |
| Low | Eliminate render-blocking resources FCP LCP | Potential savings of 67ms |
| Low | Avoid chaining critical requests FCP LCP | 5 chains found |
| Low | Avoid enormous network payloads LCP | Total size was 698KB |
| Low | Reduce unused CSS FCP LCP | Potential savings of 44.3KB |
| Low | Reduce initial server response time FCP LCP | Root document took 125ms |
| Low | Serve static assets with an efficient cache policy | Potential savings of 7.25KB |
| Low | Serve images in next-gen formats | Potential savings of 68.7KB |
| Low | Properly size images | Potential savings of 116KB |
| Low | Defer offscreen images | Potential savings of 208KB |
| N/A | Largest Contentful Paint element LCP | 440 ms |
| N/A | Reduce JavaScript execution time TBT | 9ms spent executing JavaScript |
| N/A | Avoid large layout shifts CLS | 4 elements found |
| N/A | Reduce the impact of third-party code TBT | Total size was 310KB |
| N/A | Minimize main-thread work TBT | Main-thread busy for 187ms |
| N/A | Avoid serving legacy JavaScript to modern browsers TBT | Potential savings of 56B |
| N/A | Avoid an excessive DOM size TBT | 72 elements |
| N/A | User Timing marks and measures | |
